# Bandpass corticospinal transmission during repetitive TMS revealed by motor unit recordings

**DOI:** 10.64898/2026.05.20.726653

**Authors:** Hélio V. Cabral, Milena A. Dos Santos, Andrea Rizzardi, J. Greig Inglis, Maria Cristina Rizzetti, Andrea Pilotto, Alessandro Padovani, Francesco Negro

## Abstract

We employed a noninvasive high-density surface electromyography (HDsEMG) framework to track spinal motor neuron responses during repetitive transcranial magnetic stimulation (rTMS) and characterize the transmission of rhythmic corticospinal inputs to the alpha motor neuron pool. Eleven healthy individuals performed isometric thumb flexion while rTMS was delivered over the motor cortex at five frequencies (5, 10, 20, 30, and 50 Hz) and three subthreshold intensities (50%, 60%, and 70% of resting motor threshold). Motor units were decomposed from HDsEMG before stimulation and tracked during rTMS, allowing input-output coupling between the stimulation train and individual motor unit and cumulative spike train activity to be quantified. rTMS inputs were significantly transmitted at all frequencies except 5 Hz, indicating widespread corticospinal coupling. At the motor neuron population level, corticospinal transmission preserved the spectral components of the rTMS input and exhibited intensity-dependent increases in coupling strength, consistent with approximately linear transmission. The estimated corticospinal transmission exhibited a bandpass-like profile, with transmission between 20 and 50 Hz significantly greater than at lower and higher frequency ranges. Transmitted rTMS inputs induced frequency-specific increases in the common synaptic input to motor neurons, with oscillations at the stimulation frequency significantly enhanced during rTMS compared with before stimulation. Computational simulations reproduced this frequency selectivity only when excitatory and inhibitory postsynaptic inputs were combined. These findings reveal frequency-selective, approximately linear corticospinal transmission during rTMS and establish a noninvasive framework for assessing corticospinal function, with potential application as a tool for tracking changes associated with neurological disease and response to neurorehabilitation.

## I. INTRODUCTION

It is well established that the corticospinal tract plays an essential role in the control of skilled human movement, particularly in actions involving the upper limb and hand muscles [1, 2]. Evidence from human and non-human primates shows that lesions to the primary motor cortex [3–6] or corticospinal tract [7–9] lead to marked impairments in dexterous hand movements. These deficits are closely linked to the presence of direct cortico-motoneuronal connections, which provide monosynaptic projections from neurons in the primary motor cortex to spinal motor neurons [2]. Such projections are more prominent in species with a greater capacity for fine motor control [10–12] and are particularly strong for distal muscles, including those controlling the fingers [13, 14]. Consistent with this, even weaker cortical stimulation can evoke large excitatory post-synaptic potentials in hand muscles, reflecting the high efficacy of corticospinal transmission [14, 15]. Together, these findings highlight monosynaptic corticospinal projections as a key element for the precise control of voluntary movement.

Electrophysiological studies combining cortical and muscle recordings further support the central role of shared synaptic input to spinal motor neurons in voluntary human movement. Oscillatory activity in the cortex, measured using electroencephalography or magnetoencephalography, shows significant coupling with oscillations in surface electromyograms (EMG) or motor unit spike trains from the contralateral muscle [16–21]. These corticomuscular interactions, commonly referred as to corticomuscular coherence, are most consistently observed in the ∼15-35 Hz frequency range (beta band), suggesting that corticospinal pathways preferentially transmit neural oscillations within this band during sustained contractions [16]. However, studies of motor unit activity indicate that the bandwidth of shared synaptic inputs reaching spinal motor neuron pools extends to higher frequencies, up to approximately 70-80 Hz [22–25]. This discrepancy suggests that the transformation of cortical inputs into motor neuron output is not a simple transmission process but may involve frequency-dependent filtering or modulation within the corticospinal pathway [26, 27].

Understanding the input-output properties governing corticospinal transmission, such as frequency selectivity, is fundamental for comprehending how the nervous system controls precise and skilled movements. Transcranial magnetic stimulation (TMS), and its repeated variant (repetitive TMS; rTMS), provide a non-invasive way to assess these properties by delivering controlled inputs to the motor cortex [28]. TMS uses an electromagnetic coil placed over the scalp to generate a magnetic field, which induces weak electric currents in cortical neurons and can elicit motor-evoked potentials (MEPs) in contralateral muscles (for a review, see Hallett [29]). These responses are commonly interpreted as reflecting the activation of corticospinal projections and are widely used as markers of corticospinal integrity and transmission efficacy. Notably, TMS-based neuromodulation protocols, combined or not with peripheral stimulation inputs (i.e., paired associative stimulation), have been shown to induce plastic changes at cortico-motoneuronal synapses and enhance motor recovery [30, 31]. Previous studies combining TMS with muscle recordings have also provided insights into the input-output relationship of the corticospinal pathway [32–35]. In particular, increasing stimulation intensity leads to larger MEP amplitudes, following a sigmoidal input-output relationship [36, 37], and to a higher probability of motor unit discharge [32, 33]. In addition, rTMS protocols have demonstrated frequency-dependent modulation of corticospinal excitability, with low-frequency stimulation typically producing inhibitory effects and high-frequency stimulation producing facilitatory effects, as reflected by changes in MEP amplitude [38–40]. However, despite these advances, how rTMS-induced oscillatory inputs at different frequencies are transmitted through the corticospinal pathway to spinal motor neurons remains poorly understood. Moreover, most studies have relied on global surface EMG or selective intramuscular recordings, which limit the characterization of corticospinal input-output transformations.

In this study, we developed a noninvasive high-density surface EMG (HDsEMG) framework to track spinal motor neuron activity during rTMS and to characterize how inputs of different frequencies and intensities are transmitted through corticospinal pathways in healthy individuals. We specifically tested whether rTMS-induced oscillatory inputs are conveyed to the motor neuron pool of hand muscles in an approximately linear manner, and whether corticospinal transmission exhibits frequency selectivity. We found that corticospinal pathways act as a bandpass system, preferentially transmitting higher-frequency components to the neural drive. By combining experimental measurements with computational modeling, we further identified potential synaptic mechanisms underlying this frequency-dependent transmission. Together, these findings provide a framework for understanding corticospinal input-output transformations and offer new insight into how descending neural signals are converted into motor commands.

## II. METHODS

### A. Participants

Eleven healthy volunteers (3 males, age 29 ± 3 years) participated in the study. All participants were right-handed, had no history of neurological or musculoskeletal disorders, and provided written informed consent before participation. Participants were recruited among university personnel and patients’ caregivers. Exclusion criteria included any contraindication to TMS, such as a history of epilepsy or seizures, metallic or electronic implants in the head or neck, cardiac pacemakers, or pregnancy. All volunteers completed a standardized TMS safety screening questionnaire prior to testing to confirm eligibility. The experimental procedures were approved by the local ethics committee (number NP1471) and conducted in accordance with the Declaration of Helsinki.

### B. Experimental Protocol

The study consisted of a single experimental session lasting approximately two hours. Participants were seated comfortably in a chair with their right forearm positioned on a custom-built support secured to the testing table (**Fig. 1A**, left). The elbow and wrist were positioned at approximately 45° of flexion and midway between full supination and full pronation, respectively. The four fingers were aligned with the forearm, and the thumb was maintained in a mid-flexed position. The proximal segment of thumb was attached to an adjustable support connected to a load cell (SM-100 N; Interface, Arizona, USA) used to record isometric thumb flexion force (**Fig. 1A**, top left). The forearm, wrist and four fingers were firmly stabilized with Velcro straps to minimize movement.

**Fig. 1.**
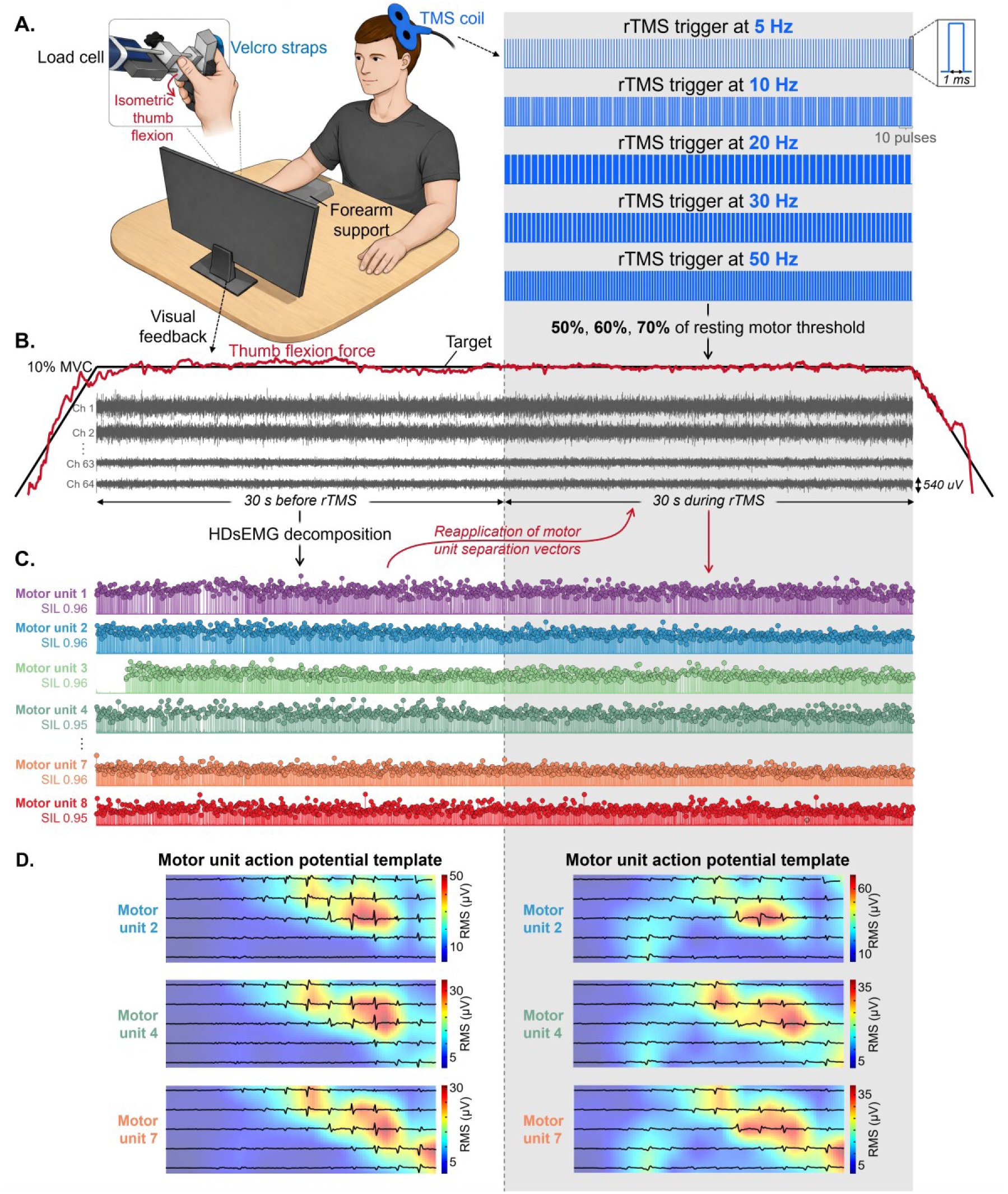
Experimental framework to assess spinal motor neuron activity during rTMS. (A) High-density surface electromyograms (HDsEMG) were recorded from the thenar muscles using a 64-channel grid while participants performed isometric thumb flexion at 10% of maximal voluntary contraction (MVC). The thumb was attached to a load cell to measure force, and participants received real-time visual feedback of force output. Repetitive transcranial magnetic stimulation (rTMS) was delivered with different frequencies (5-50 Hz) and intensities (50-70% resting motor threshold; RMT), in randomized order. (B) Representative HDsEMG signals from selected channels during a 60 s contraction, including 30 s before and 30 s during rTMS. The force signal (red) closely matched the target (black) in both periods. (C) Motor unit decomposition and tracking. Motor unit spike trains were identified from HDsEMG signals recorded before rTMS using convolutive blind source separation (left) and tracked during rTMS by reapplying the corresponding separation vectors (right). Each color represents a different motor unit. (D) Representative two-dimensional motor unit action potential (MUAP) templates for tracked motor units before and during rTMS, demonstrating high spatial similarity and confirming successful tracking across conditions.

After RMT identification (see next section), participants performed submaximal isometric thumb flexion tasks both before and during rTMS. Initially, three maximal voluntary contractions (MVCs) of isometric thumb flexion were performed for 5 s each, separated by a 3-min rest interval. The highest peak force across trials was defined as the MVC and used to set the target for subsequent submaximal tasks. Participants then performed isometric thumb flexion at 10% MVC. Each trial consisted of a ramp-up phase from 0% to 10% MVC at 5% MVC/s, a steady plateau phase lasting 60 s, and a ramp-down phase from 10% to 0% MVC at - 5% MVC/s. During the plateau phase, participants maintained steady contraction for 30 s before and 30 s during rTMS (**Fig. 1B**). Five stimulation frequencies and three intensities were delivered, yielding 15 trials per participant. At least a 3-min rest period was provided between trials. HDsEMG signals were recorded from thenar muscles simultaneously with thumb flexion force throughout all trials.

### C. Resting Motor Threshold Identification and rTMS Protocol

TMS was delivered with a figure-of-eight coil connected to a DuoMAG rTMS xt-100 (DEYMED Diagnostic s.r.o, Czech Republic). The coil was held tangentially to the scalp over the left primary motor cortex with the handle pointing posterior-laterally at approximately 45° to the sagittal midline, inducing a posterior-anterior current in the motor cortex [41, 42]. Initially, the optimal stimulation site (“hotspot”), defined as the spot on the scalp where the largest MEPs in the thenar muscles were elicited at the lowest TMS intensity, was identified. The hotspot was marked on the scalp using a neuronavigation system (SofTaxic Neuronagivation System, v 3.0, E.M.S., Bologna, Italy), which was subsequently used throughout the experiment to ensure consistent coil positioning.

The RMT was determined as the minimum stimulation intensity that evoked MEP responses equal or greater than 50 μV in the thenar muscles in at least five out of ten consecutive stimulations while the muscle was at rest (Rossini-Rothwell method) [28, 43]. During RMT identification, single 290-µs biphasic pulses were delivered at 0.1 Hz. RMT was used as the reference for determining stimulation intensity to provide an individually normalized measure without requiring participants to maintain a voluntary contraction during threshold identification, thereby minimizing additional muscle activation and potential fatigue during the experimental session. Individual RMT values were 41%, 43%, 36%, 41%, 38%, 40%, 41%, 41%, 45%, 39%, and 41% of maximal stimulator output (MSO), with a group average of 40.5 ± 2.4% MSO. Once identified, the RMT was used as the reference to set subthreshold stimulation intensities for the submaximal isometric tasks.

Five stimulation frequencies (5, 10, 20, 30, and 50 Hz) and three stimulation intensities (50%, 60%, and 70% RMT) were delivered in a randomized order, resulting in a total of 15 stimulation protocols. The absolute stimulation intensities corresponding to 50%, 60%, and 70% RMT were, on average, 20.3 ± 1.2%, 24.3 ± 1.4%, and 28.4 ± 1.7% MSO, respectively. Stimulation included 5 Hz and higher frequencies, as rTMS at or above 5 Hz has been reported to increase corticospinal excitability [28, 40]. Submotor-threshold intensities were selected to avoid evoking overt motor responses and associated sensory reafference during the voluntary submaximal contraction. For each frequency-intensity condition, rTMS was delivered over a 30 s period using biphasic magnetic pulses with a duration of 290 µs. At 5 Hz, pulses were delivered continuously at 200 ms intervals, corresponding to a total of 150 pulses. For stimulation frequencies above 5 Hz (10, 20, 30, and 50 Hz), pulses were delivered in blocks of 10, with successive pulses within each block separated by intervals determined by the stimulation frequency (100, 50, 33.3, and 20 ms, respectively). Consecutive blocks were separated by 200 ms intervals. Thus, although the total stimulation period was matched across frequencies (30 s), the number of delivered pulses and their temporal distribution varied across conditions.

### D. HDsEMG and Force Recordings

HDsEMG signals were recorded from the thenar muscles using a 64-channel electrode grid (13 rows × 5 columns, 4 mm interelectrode distance; HD04MM1305, OT Bioelettronica, Turin, Italy). **Fig. 1B** shows representative HDsEMG signals acquired during the 60 s contraction. Before electrode placement, the skin region was shaved (when necessary) and cleaned with abrasive paste (EVERI, Spes Medica, Genova, Italy) and water to reduce impedance. The grid was fixed to the skin using bi-adhesive foam, and electrode-skin contact was ensured by filling the foam cavities with conductive paste (AC Cream, Spes Medica, Genova, Italy). A strap reference electrode was positioned over the right wrist.

HDsEMG signals were recorded in monopolar mode and digitized at 2,000 Hz using a 16-bits analog to digital converter (Sessantaquattro+, OT Bioelettronica, Turin, Italy) with a bandwidth of 10-500 Hz. Thumb flexion force was measured using a load cell (SM-100 N; Interface, Arizona, USA) and amplified by a factor of 100 (ForzaJ, OT Bioelettronica, Turin, Italy). The timing of each stimulation pulse was recorded using a 1 ms rectangular trigger signal, which was used to construct the rTMS input trains for the analyses (**Fig. 1A**, right). Both force and trigger signals were acquired as auxiliary channels in the HDsEMG system. Participants received real-time visual feedback of the target force and their produced force on a computer monitor to maintain the required force level during the task (**Fig. 1A**, left).

### E. Simulations

To explore mechanisms underlying the frequency selectivity observed experimentally (see Results), we simulated motor neuron pool responses to different synaptic input scenarios resulting from rTMS. Simulations were implemented using the motor neuron pool model proposed by Fuglevand, et al. [44]. Detailed descriptions of this modeling framework can be found in previous studies [45, 46]. Motor neuron parameters were consistent with those reported by Cisi and Kohn [47], and the number of simulated motor neurons was set to six to match the average number of motor units tracked experimentally.

The input to the motor neuron pool was modeled as the linear summation of a common synaptic input shared across all motor neurons and an independent noise input specific to each motor neuron [45]. The independent input, representing variability in the membrane potential of individual motor neurons, was modeled as Gaussian noise with a bandwidth of 50 Hz. The common synaptic input was modeled as a train of post-synaptic potentials (PSPs) triggered by rTMS inputs. Although individual TMS pulses may evoke multiple descending corticospinal volleys (i.e., D- and I-waves) [48], resulting in temporally summated PSPs at spinal motor neurons, our simulations did not explicitly model these individual volleys. Instead, to simplify the model and focus on the synaptic mechanisms underlying the experimentally observed frequency selectivity, the aggregate postsynaptic response to each stimulation pulse was approximated by a single effective PSP. Three synaptic input scenarios were simulated: (1) a train of excitatory post-synaptic potentials (EPSPs) only, (2) a train of inhibitory post-synaptic potentials (IPSPs) only, and (3) a train of combined EPSP and IPSP inputs (**Fig. 6A**). For each scenario, input trains were generated at the same stimulation frequencies used experimentally (5, 10, 20, 30, and 50 Hz).

The synaptic response to each rTMS pulse was modeled using an alpha function kernel (equation (1)), a common approximation of synaptic dynamics in computational models [49, 50].

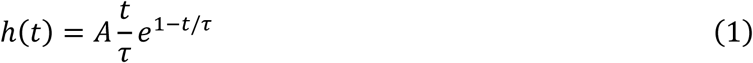

where *A* represents the amplitude (gain) of the post-synaptic potential and *τ* defines the temporal dynamics of the synaptic response, rather than its total duration. For each stimulation frequency, the input train was convolved with an EPSP kernel, an IPSP kernel, or a combination of both kernels depending on the simulated scenario. For EPSPs the gain parameter *A* was positive, whereas for IPSPs it was negative.

Different synaptic parameters were explored to determine which combination best reproduced the experimentally observed frequency-dependent transmission (see Results). Excitatory synaptic responses were modeled with time constants *τ* = 2 or 4 ms to represent relatively fast postsynaptic dynamics, consistent with electrophysiological observations in animal models of AMPA receptor-mediated transmission [51]. These values are also compatible with indirect estimates from human experiments indicating that corticospinal excitatory postsynaptic potentials can persist for approximately 10 ms following descending volleys [52]. Inhibitory synaptic responses were modeled using longer time constants (*τ* = 6 or 10 ms) to represent relatively slower postsynaptic dynamics, consistent with the time course of glycinergic-mediated transmission observed in motor neurons [53, 54]. Two gain values were simulated (*A* = 0.0007 and *A* = 0.0015). This allowed the combined EPSP-IPSP scenario to generate three relative gain configurations: equal gain for EPSPs and IPSPs, EPSP gain twice that of IPSPs, and IPSP gain twice that of EPSPs. These gain values were selected after preliminary testing to approximate the range of coupling values observed experimentally.

All combinations of synaptic time constants and gains were tested, resulting in four parameter combinations for the EPSP-only scenario, four for the IPSP-only scenario, and sixteen for the combined EPSP–IPSP scenario. In the combined scenario, the IPSP was delayed relative to the EPSP by 2 ms, representing a feedforward inhibitory mechanism. For each simulated condition, the spectral input-output coupling profile of corticospinal transmission was estimated using the same analysis procedures applied to the experimental data (see Data Analysis).

### F. Data Analysis

#### 1 Force Analysis

Force signals were low-pass filtered at 15 Hz using a third-order Butterworth filter. To quantify force steadiness, we calculated the coefficient of variation of the detrended force signal (standard deviation divided by the mean) over the 30 s period before rTMS and the 30 s period during rTMS.

#### 2 Bipolar EMG Amplitude

To compare EMG amplitude before and during rTMS, we calculated the root-mean-square (RMS) amplitude of a simulated bipolar EMG channel located at the center of the electrode grid. The bipolar signal was obtained by subtracting the average of six monopolar channels located on rows 8 and 9 (columns 1-3) from the average of six monopolar channels located on rows 5 and 6 (columns 1-3). The RMS amplitude was computed over non-overlapping windows of 250 ms across the 30 s epochs before and during rTMS. The RMS values obtained from all windows were then averaged to obtain a single value for each epoch. To allow comparison across participants, RMS amplitude was normalized to the RMS value obtained during the MVC. This MVC RMS value was calculated from a 250 ms window centered on the peak force during the MVC trial.

#### 3 Motor Unit Decomposition

HDsEMG signals were first band-pass filtered with a third-order Butterworth filter (20-500 Hz cutoff frequencies). Channels with low signal-to-noise ratio or visible artifacts were identified through visual inspection and discarded from further analysis. Motor unit discharge times were then extracted from the HDsEMG signals recorded during the 30 s period preceding rTMS using a convolutive blind source separation algorithm [55]. The decomposition was performed by one of the authors (H.V.C.) using an implementation of the algorithm developed by another author of this study (F.N.). This method has been previously validated and widely applied to extract single motor unit activity from HDsEMG recordings of hand muscles [55, 56]. Briefly, the HDsEMG signals were first temporally extended by adding delayed versions of each channel and subsequently whitened to decorrelate the extended observations. An iterative source extraction procedure based on a fast independent component analysis (fastICA) algorithm was then applied to identify individual motor unit sources. During this approach, a contrast function was optimized to maximize the sparsity of the estimated sources (i.e., motor unit innervation pulse trains, IPTs) and identify the separation vectors of each unit [55, 57]. Each separation vector defines a unique spatiotemporal filter that, when convolved with the HDsEMG signals, provides an estimate of the corresponding motor unit IPT. Discharge times were then identified from the IPTs through peak detection followed by *k-means* clustering into two groups, separating motor unit discharges from background noise activity. Thereafter, the initial source estimates were further refined through an iterative procedure that seeks to minimize the discharge rate variability. Representative motor unit IPTs are shown in **Fig. 1C** (left), with each unit represented in a different color and circles indicating the discharge times. Further details of the decomposition algorithm are provided in the original study [55]. Only motor units with a silhouette value greater than 0.90 were retained for further analysis. The silhouette value quantifies the separation between the clusters corresponding to motor unit discharges and background activity in the estimated IPTs and is commonly used as an indicator of decomposition quality [55]. However, because a high SIL value alone does not guarantee reliable identification of all discharge times [58], the decomposed motor unit spike trains were subsequently visually inspected by two expert operators (H.V.C. and M.A.S.) to identify false positives and false negatives. Missing or incorrectly identified discharges producing nonphysiological discharge rates were manually corrected, and the motor unit spike trains were iteratively re-estimated as previously described [59]. This manual editing procedure has been shown to produce highly reliable results across operators [60]. We also attempted to independently decompose the HDsEMG signals recorded during rTMS, but this approach resulted in a low yield and poor decomposition quality, even considering that subthreshold stimulation intensities were used.

#### 4 Motor Unit Tracking

To track motor units across time periods (before- and during-rTMS), the separation vectors obtained from the blind source separation algorithm during the pre-rTMS period were reapplied to the HDsEMG signals recorded during rTMS [56, 61, 62], allowing reliable identification of the same motor units across time. Importantly, motor unit tracking was performed independently for each frequency-intensity condition, with the same motor units identified before and during rTMS within each condition, but without attempting to track them across different stimulation frequencies or intensities. To verify successful tracking, we calculated the two-dimensional motor unit action potential (MUAP) templates using the spike-triggered averaging technique [59, 63] and their root-mean-square amplitude. **Fig. 1D** shows three examples of MUAP templates for tracked units, demonstrating a high degree of similarity in MUAP shape and amplitude between time periods and thereby confirming that the same units were successfully tracked.

#### 5 Input-output Coherence Analysis

To assess whether rTMS inputs were commonly transmitted to spinal motor neurons of the thenar muscles, input-output coupling was quantified by computing the coherence between the rTMS train (input) and individual motor unit spike trains (outputs). The rTMS train was represented as a binary time series marking the timing of each stimulation pulse, identified from the recorded 1 ms rectangular trigger signal. Coherence is a frequency-domain measure that quantifies the linear association between two signals as a function of frequency and ranges from 0 (no coupling) to 1 (perfect coupling). The rTMS train was represented as a binary time series marking the timing of each stimulation pulse. Motor unit discharge times were also converted to binary spike trains. Coherence between the rTMS train and each motor unit spike train was estimated using the Welch’s periodogram method with a 1 s Hanning window and an overlap of 950 ms [64]. To determine the significance of the observed coupling, coherence was also calculated after randomly shuffling the interspike intervals of each motor unit spike train while preserving its firing statistics. This procedure generated surrogate spike trains with identical discharge rate distributions but without temporal coupling to the rTMS input. For each rTMS frequency, the strength of input-output coupling was quantified as the area under the coherence curve within a ± 1 Hz window centered on the stimulation frequency. This metric was used to compare coupling strength across stimulation frequencies and conditions. Because the strongest input-output coupling was observed at 70% RMT (see Results), the input-output coherence analysis was restricted to this stimulation intensity.

#### 6 Assessment of Approximately Linear Corticospinal Transmission

To test whether corticospinal transmission during rTMS exhibited properties consistent with approximately linear transmission at the motor neuron population level, coherence was calculated between the rTMS train and the cumulative spike train (CST) of all identified motor units. The CST was obtained by summing the binary spike trains of all motor units identified in each condition and represents the neural drive to the muscle. Because coherence estimates are influenced by the number of spike trains contributing to the CST [22], the minimal number of motor units identified across rTMS stimulation intensities within each participant was used to construct the CST. When more motor units were available in a given condition, the CST was computed by randomly selecting the minimal number of motor units from the available pool. This procedure was repeated up to 100 permutations, and the average coherence across all iterations was used for further analysis. Coherence between the rTMS train and the CST was estimated using Welch’s periodogram with a 1 s Hanning window and an overlap of 950 ms. To stabilize variance and allow comparisons across conditions, coherence values were transformed in z-scores as described in Gallet and Julien [65], which accounts for the bias introduced by overlapping windows. To assess the preservation of input frequency components, we examined whether the CST spectrum reproduced the frequencies present in the rTMS input. Specifically, the strength of rTMS-CST coupling was quantified as the area under the coherence curve within ± 1 Hz around the stimulation frequency. To determine statistical significance, coherence was also computed using surrogate CST signals generated by shuffling the interspike intervals of the motor unit spike trains while preserving their firing statistics. Because the strongest input-output coupling was observed at 70% RMT (see Results), the analysis was restricted to this stimulation intensity. To assess the intensity dependence of input-output coupling, we evaluated whether rTMS-CST coupling increased proportionally with stimulation intensity. For each stimulation frequency, the relationship between stimulation intensity (50%, 60%, and 70% RMT) and rTMS-CST coupling at the stimulation frequency was assessed using linear regression.

#### 7 Spectral Input-output Coupling Profile

To characterize the frequency-dependent transmission properties of the corticospinal system during rTMS, we estimated a spectral input-output coupling profile. For each stimulation frequency, the area under the curve of the rTMS-CST z-coherence was calculated within a ± 1 Hz window around the frequencies of interest (5, 10, 20, 30, 40, 50, 60, 70, and 80 Hz), corresponding to the fundamental stimulation frequencies and their harmonic components. Importantly, frequencies above 50 Hz were analyzed exclusively as harmonics of the stimulation trains and were not independently delivered as stimulation frequencies. The spectral input-output coupling profile was then estimated as the ratio between the area under the curve obtained from the original CST and the area under the curve obtained from the shuffled CST. To facilitate interpretation, we subtracted 1 from the resulting ratio. Thus, values greater than 0 indicated an increase in rTMS-CST coupling relative to the shuffled condition, whereas values close to 0 indicated no difference.

#### 8 Motor Unit Pooled Coherence

To assess changes in common synaptic input to spinal motor neurons during rTMS, coherence analysis was performed between motor unit spike trains [22, 62, 64, 66]. Coherence was calculated between two CSTs, each obtained by summing the binary spike trains of a subset of identified motor units. Each CST specifically contained half of the available motor unit spike trains. Only participants with at least four identified motor units were included in this analysis (N = 7). Coherence was computed between the two detrended CSTs using Welch’s periodogram method with a 1-s Hanning window and 950 ms overlap. This procedure was repeated for up to 100 random permutations of motor unit grouping, and the average coherence across all permutations was calculated to obtain the pooled coherence. The pooled coherence was transformed into z-scores [65]. To determine the confidence level for significant coherence, the bias was estimated as the mean z-coherence value between 250 and 500 Hz, a frequency range where no physiological coherence is expected [22]. To quantify changes in common oscillatory input associated with rTMS, the strength of motor unit-motor unit coupling was expressed as the area under the curve of the z-coherence spectrum within ± 1 Hz around the rTMS stimulation frequency.

### G. Statistical Analysis

All statistical analyses were performed in R (version 4.4.2, R Foundation for Statistical Computing, Vienna, Austria), using the RStudio environment.

To compare rTMS-motor unit spike train coupling at the main stimulation frequency between non-shuffled and shuffled spike trains (**Fig. 2B**), linear mixed-effects models (LMMs) were applied. The models included spike train type (non-shuffled and shuffled), rTMS frequency (5, 10, 20, 30, and 50 Hz), and their interaction as fixed effects, and participant as a random effect (random intercept model). Including participant as a random intercept accounted for the non-independence of motor unit observations clustered within the same participant, thereby avoiding the assumption that motor units recorded from the same participant represented independent experimental subjects. The same statistical approach was used to compare rTMS-CST coupling at the main stimulation frequency (**Fig. 3B**). LMMs were implemented using the *lmerTest* package [67]. The Kenward-Roger method was used to approximate the degrees of freedom and estimate P-values. When significant effects were detected, pairwise comparisons were performed using the *emmeans* package to obtain estimated marginal means and their differences with 95% confidence intervals [68]. Pairwise contrasts were computed separately for each rTMS frequency, and additional contrast analyses (difference-of-differences) were performed when appropriate to compare changes between shuffled and non-shuffled spike trains across rTMS stimulation frequencies.

**Fig. 2.**
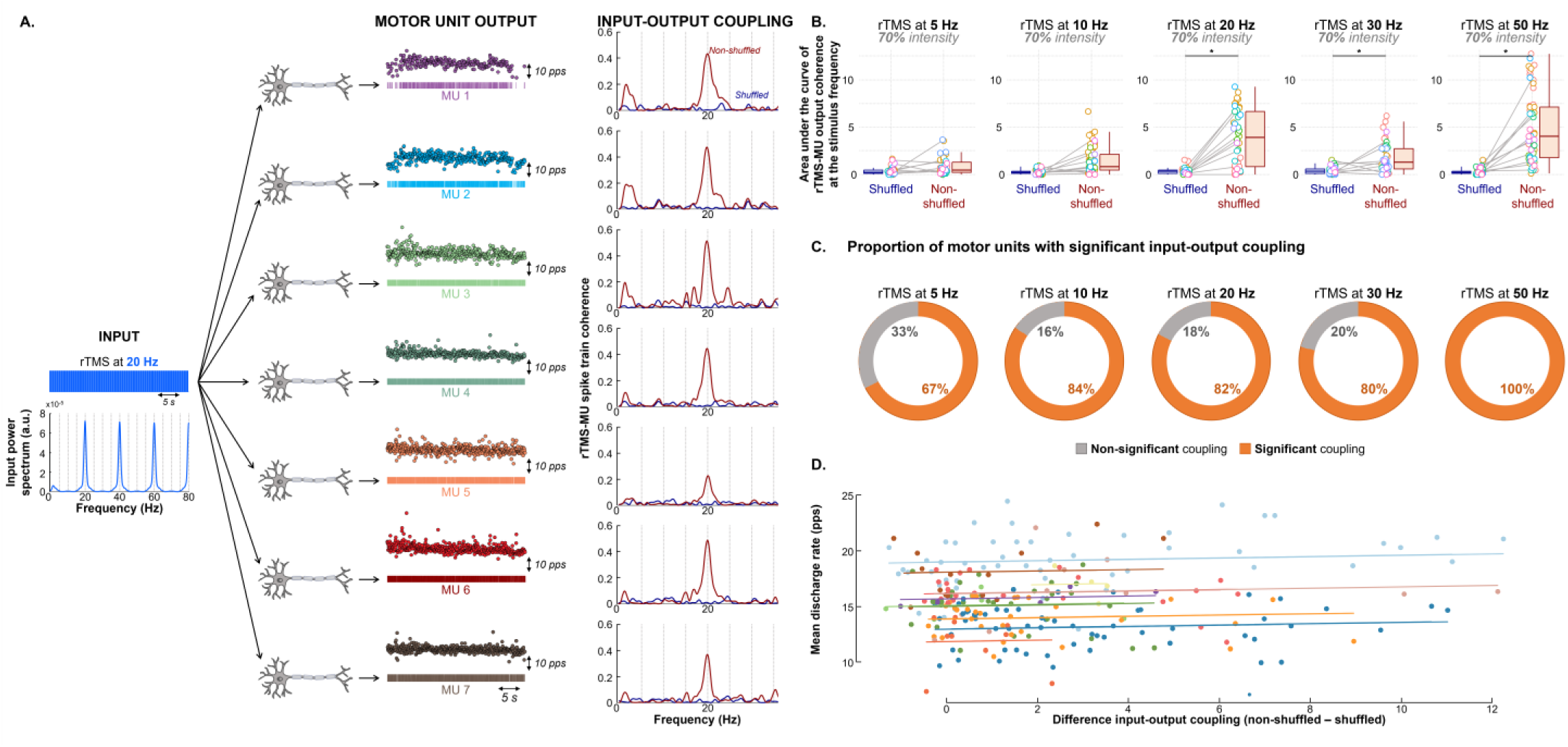
Transmission of rTMS oscillations to individual spinal motor neurons. (A) Representative example of input-output coherence between the rTMS train (input) and individual motor unit spike trains (output) at 20 Hz stimulation (70% of resting motor threshold). The rTMS spectrum shows a peak at stimulation frequency and its harmonics. Coherence between rTMS and motor unit spike trains (dark red) shows a peak at the stimulation frequency, which is absent when coherence was calculated using the shuffled spike train (dark blue). (B) Group results showing the area under the curve of input-output coherence at the main stimulation frequency for each rTMS frequency. * indicates that coherence was significantly greater in the non-shuffled than shuffled. (C) Percentage of motor units exhibiting significant input-output coupling across frequencies. For frequencies above 5 Hz, more than 80% of motor units showed significant coupling. (D) Relationship between input-output coupling (non-shuffled minus shuffled) and motor unit discharge rate, assessed using repeated-measures correlation analysis. Each circle represents an individual motor unit, and different colors indicate different participants. No significant association was observed (R = 0.08, 95% CI: −0.05 to 0.20, P = 0.300).

**Fig. 3.**
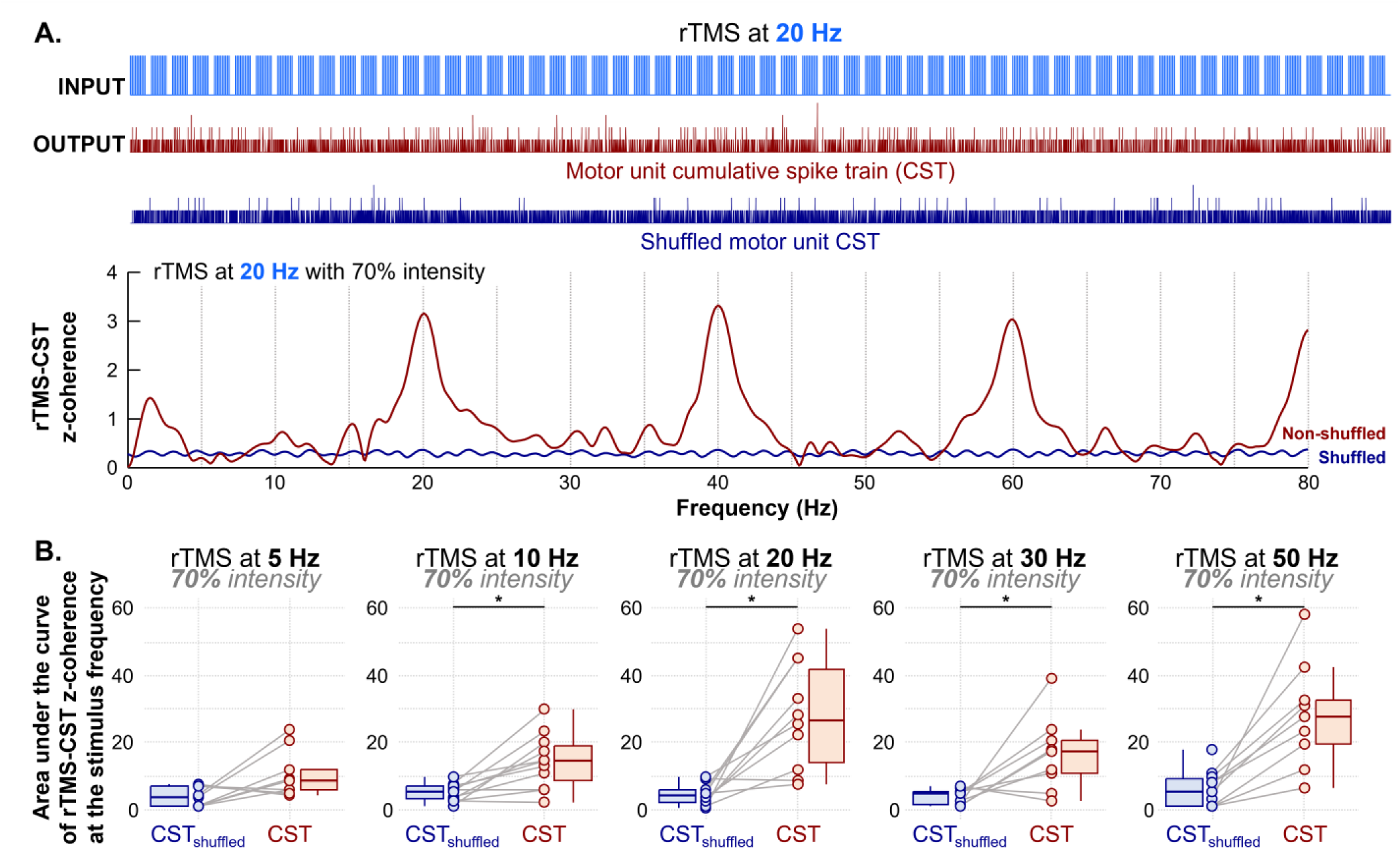
Corticospinal transmission preserved the spectral components of the rTMS input. (A) Representative example of z-coherence between the rTMS train and the cumulative spike train (CST) at 20 Hz stimulation (70% of resting motor threshold). Peaks at the stimulation frequency and its harmonics are present in the rTMS-CST coherence (dark red) but absent in the shuffled CST (dark blue), indicating that output spectra reproduce input frequencies. (B) Group results showing z-coherence at the main stimulation frequency. * indicates significant increases in input-output coupling.

To quantify how consistently rTMS inputs were transmitted across motor neurons, we calculated the percentage of motor units showing greater coupling with the rTMS input than with the shuffled spike trains (**Fig. 2C**). To determine whether differences in rTMS-motor unit spike train coupling between non-shuffled and shuffled spike trains were associated with motor unit mean discharge rate, repeated-measures correlation analysis was performed (**Fig. 2D**).

For the spectral input-output coupling profile analysis, the area-under-the-curve ratios were tested against zero using one-sample Wilcoxon signed-rank tests (null hypothesis µ_0_ = 0), performed separately for each frequency. The effects of coherence frequency and rTMS intensity on the ratios were further assessed using a linear mixed-effects model, including frequency, rTMS intensity, and their interaction as fixed effects and participant as a random effect. To characterize the frequency-dependent transmission profile (**Fig. 4C**), contrasts of estimated marginal means compared the ratios across low (5-10 Hz), intermediate (20-50 Hz), and high (60-80 Hz) frequency ranges.

**Fig. 4.**
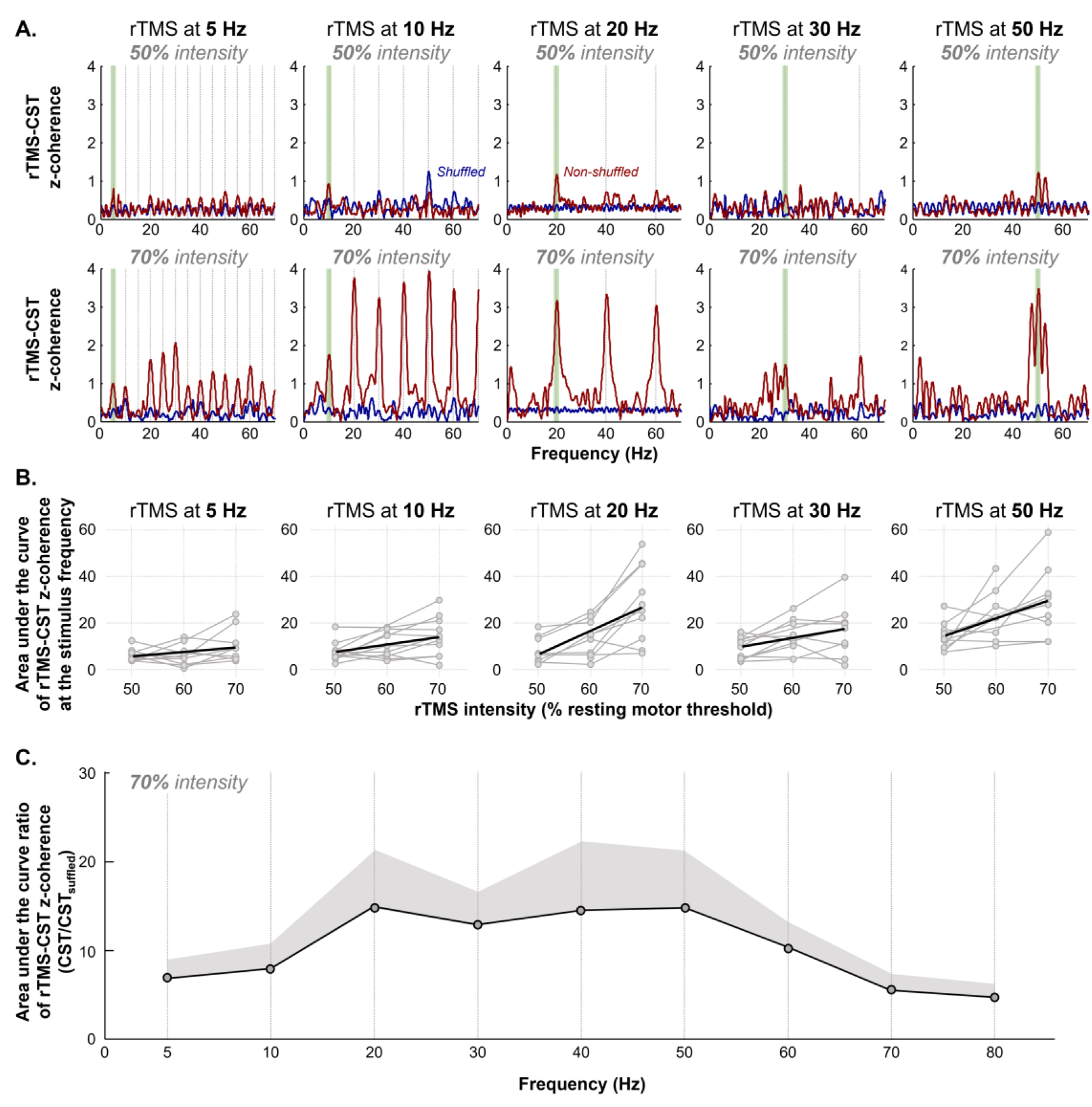
Corticospinal transmission scales with stimulation intensity and exhibits a bandpass-like behavior. (A) Representative examples of rTMS-CST z-coherence at 50% and 70% of resting motor threshold across stimulation frequencies. Input-output coupling at the stimulation frequency (green rectangles) increases with intensity for all frequencies above 5 Hz. (B) Group results showing the relationship between stimulation intensity and input-output coupling. A significant linear increase in coupling with intensity was observed for all frequencies except 5 Hz. (C) Group results of the spectral input-output coupling profile estimated as the ratio between rTMS-CST z-coherence and shuffled CST across frequencies (5–80 Hz). Values greater than zero indicate increased input-output coupling relative to shuffled data. Significant coupling was observed for all frequencies, except 5 Hz, with maximal transmission between 10 and 60 Hz, indicating a bandpass-like transmission of corticospinal inputs during rTMS.

To compare motor unit pooled coherence at the stimulation frequency (**Fig. 5B**), LMMs were applied with time (before rTMS vs during rTMS) and rTMS frequency as fixed effects and participant as a random effect. When significant effects were detected, pairwise comparisons and contrast analyses were performed. Statistical significance was set at P < 0.05. All individual data of motor unit discharge times during rTMS are available at https://doi.org/10.6084/m9.figshare.32129323.

**Fig. 5.**
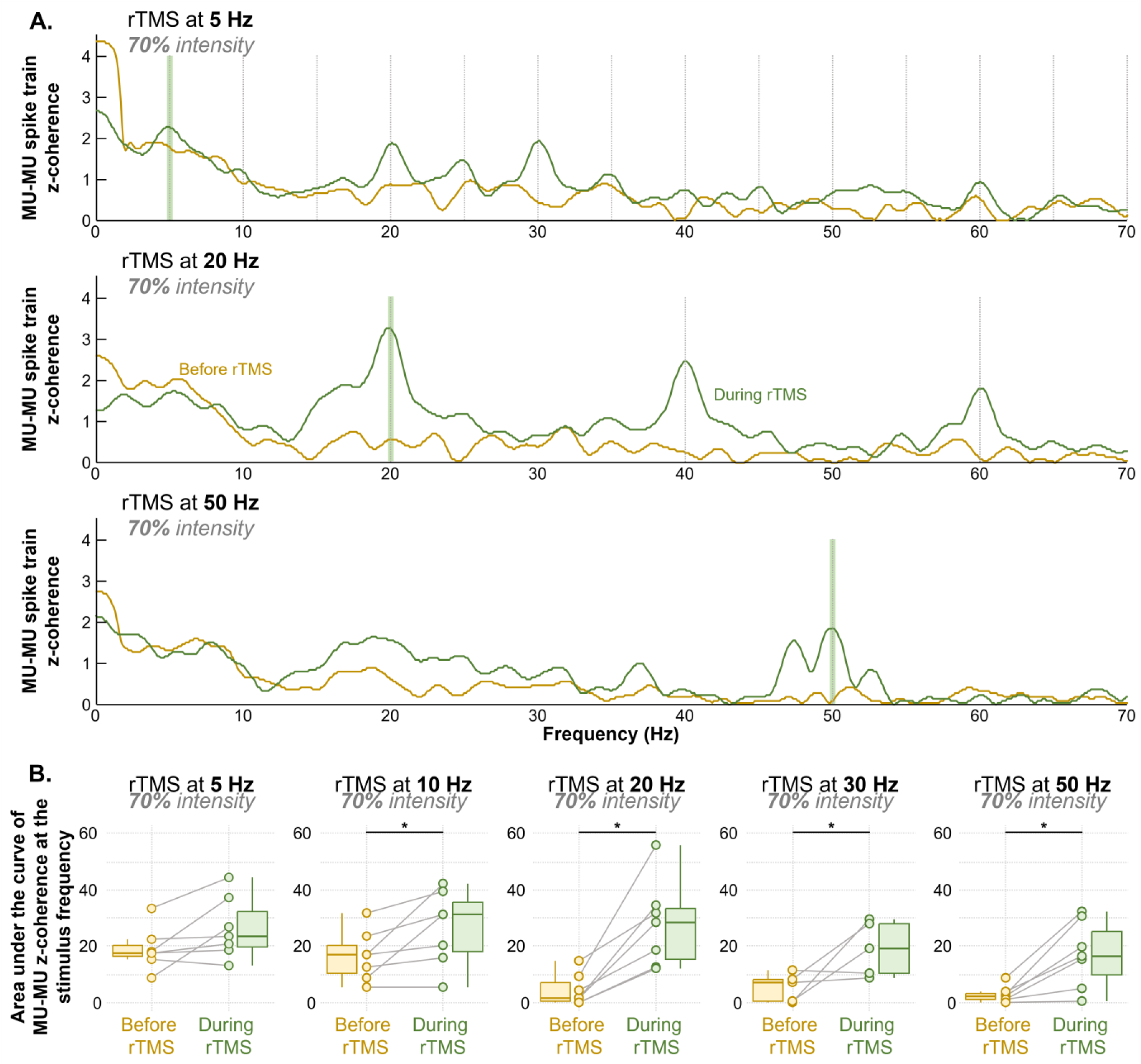
rTMS induces frequency-specific oscillations in common synaptic input. (A) Representative motor unit pooled z-coherence before (dark yellow) and during rTMS (dark green) for 5, 20, and 50 Hz stimulation. Increased coherence at the stimulation frequency (green rectangles) is evident during rTMS for 20 and 50 Hz, but not for 5 Hz. (B) Group results showing the area under the curve of motor unit pooled z-coherence at stimulation frequency. * indicates significant increases from pre-to during-rTMS.

**Fig. 6.**
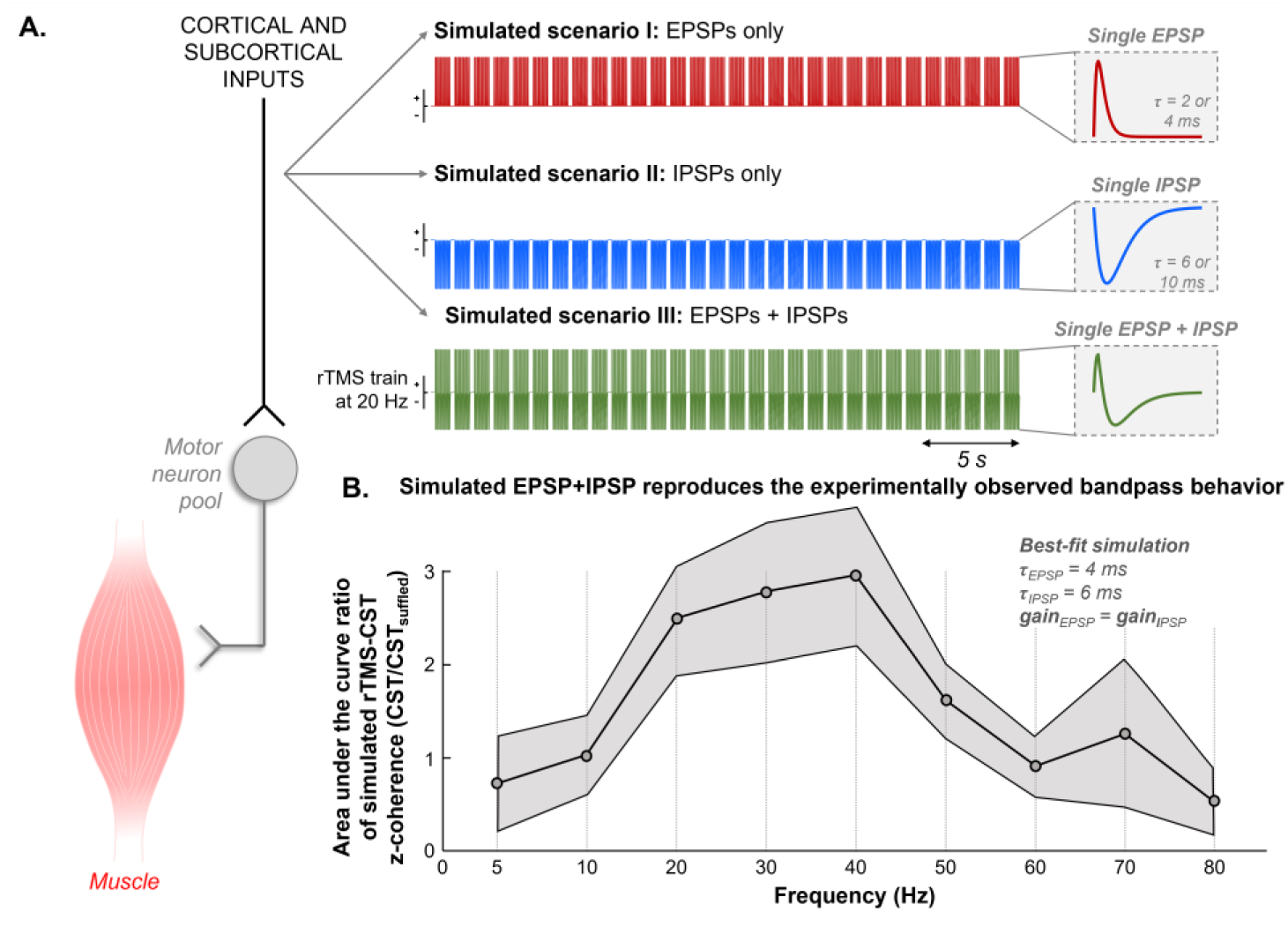
Simulated synaptic mechanisms underlying bandpass corticospinal transmission during rTMS. (A) Conceptual model of common synaptic inputs to the motor neuron pool during rTMS. Three synaptic scenarios were simulated: excitatory post-synaptic potentials (EPSPs) only, inhibitory post-synaptic potentials (IPSPs) only, and combined EPSP-IPSP inputs. Each rTMS pulse generated post-synaptic potentials modeled with an alpha function. (B) Only simulations combining excitatory and inhibitory post-synaptic potentials reproduced the bandpass filtering observed experimentally. The closest match to the experimental input-output coupling profile was obtained with τEPSP = 4 ms, τIPSP = 6 ms, and equal synaptic gain.

## III. RESULTS

### A. Subthreshold rTMS does not Alter EMG Amplitude or Force Steadiness

We used a noninvasive HDsEMG-based experimental framework to assess spinal motor neuron activity during rTMS at different stimulation frequencies and intensities. Five rTMS frequencies (5, 10, 20, 30, and 50 Hz) and three intensities (50%, 60%, and 70% of the resting motor threshold, RMT) were tested in a randomized order. Submotor-threshold stimulation was used to avoid evoking overt motor responses during the submaximal contraction. Consistent with this aim, the normalized root-mean-square amplitude for a simulated bipolar EMG at the center of the grid (%MVC) and force steadiness (coefficient of variation of force) did not significantly change between pre- and during-rTMS (Wilcoxon signed-rank test, N = 11, P > 0.193 for all comparisons). These findings confirm that the subthreshold stimulation intensities did not directly interfere with force production or EMG amplitude, ensuring that subsequent analyses reflect neural transmission. All mean values before and during rTMS are provided in **Table S1**.

### B. rTMS Oscillations are Widely Transmitted to Spinal Motor Neurons

To assess corticospinal transmission to spinal motor neurons, we decomposed motor units from HDsEMG signals recorded before rTMS and tracked their discharge activity during stimulation. An average of 12 ± 8 motor units per pre-rTMS recording were identified after decomposition and visual inspection, of which 6 ± 5 motor units were tracked between the pre- and during-rTMS periods. Having established that spinal motor unit activity could be reliably evaluated during rTMS, we next investigated whether stimulation frequencies were commonly transmitted across the motor neuron pool. To examine this, input-output coherence between the rTMS stimulation train (input) and each motor unit spike train (output) was compared with coherence obtained using a shuffled version of the same spike train, establishing a statistical confidence level for significance (**Fig. 2A**). For this analysis, we focused on 70% RMT, as this intensity consistently showed the highest input-output coupling (see next subsection). **Fig. 2A** shows an example of this analysis during 20 Hz stimulation. As expected, the rTMS train spectrum exhibited a clear peak at 20 Hz and its harmonic frequencies. For all seven motor units decomposed in this case, coherence between the rTMS train and the spike train (dark red trace) showed a distinct peak at 20 Hz that was absent when coherence was calculated using the shuffled spike train (dark blue trace).

Group analysis confirmed these observations. The area under the curve of the rTMS-motor unit output coherence at the main stimulus frequency was significantly greater than that of the shuffled version for all rTMS frequencies (linear mixed-effects models, N = 11, P < 0.026 for all), except at 5 Hz (P = 0.698) and 10 Hz (P = 0.116) (**Fig. 2B**). Across stimulation frequencies above 5 Hz, more than 80% of decomposed motor units exhibited significant coupling (**Fig. 2C**). Contrast analysis showed that increases in the rTMS-motor unit output coherence were significantly higher for 20 Hz and 50 Hz than the other stimulation frequencies (P < 0.001 for all comparisons). Importantly, differences in input-output coupling (non-shuffled minus shuffled) were not associated with mean discharge rate (**Fig. 2D**; repeated-measures correlation, R = 0.08, 95% CI [-0.05, 0.20], P = 0.300). Collectively, these results indicate that, except at 5 Hz, the rTMS input was widely transmitted to the majority of spinal motor neurons.

### C. Corticospinal Transmission during rTMS Exhibits Properties Consistent with Approximately Linear Transmission

After establishing that rTMS input is widely shared across motor neuron pool, we next asked whether corticospinal transmission exhibited properties consistent with approximately linear transmission at the motor neuron population level. First, we examined whether the CST spectrum reflected a sum of sinusoidal components at the main stimulation frequency and its harmonics. To this end, we calculated the z-coherence between the rTMS train and the CST and compared it with the z-coherence obtained using a shuffled CST (**Fig. 3A**). An example during 20 Hz stimulation is shown in **Fig. 3A**. Clear peaks at 20 Hz and its harmonics were observed in the rTMS-CST coherence (dark red), whereas these peaks were absent in the shuffled CST coherence (dark blue), indicating that the output spectrum reproduced the input frequencies. Group analysis confirmed this pattern, with significant increases in z-coherence at the main stimulus frequency for all rTMS frequencies (linear mixed-effect models, N = 11, P < 0.025 for all), except 5 Hz (P = 0.182) (**Fig. 3B**). Contrast analysis revealed that increases in z-coherence were significantly greater for 20 Hz and 50 Hz compared with the other rTMS frequencies (P < 0.025 for all comparisons).

Second, we examined whether the strength of input-output coupling increased with stimulation intensity. Representative data are shown in **Fig. 4A**, where rTMS-CST z-coherence at 50% and 70% RMT revealed larger coupling at the main stimulation frequency (green rectangles) with increasing intensity across all frequencies except 5 Hz. This intensity-dependent modulation was consistent at the group level where input-output coupling at the main stimulation frequency increased linearly with stimulation intensity for all rTMS frequencies (linear regression models, N = 11, R > 0.37, P < 0.046 for all frequency-specific regressions), except 5 Hz (R = 0.32, P = 0.076) (**Fig. 4B**). Together, these results demonstrate that, for frequencies above 5 Hz, the motor neuron population output preserves the rhythmic components of the rTMS input and exhibits intensity-dependent increases in input-output coupling, consistent with approximately linear corticospinal transmission at the motor neuron population level.

### D. Corticospinal Input-output Coupling Profile Reveals Bandpass Filtering

We next estimated the spectral input-output coupling profile of corticospinal pathways during rTMS by computing the ratio of z-coherence between the CST and shuffled CST at the stimulation frequency and its prominent harmonics (5, 10, 20, 30, 40, 50, 60, 70, and 80 Hz). A ratio near zero indicated no difference in input-output coupling between CST and shuffled CST, while values greater than zero reflected increased coupling. Group analysis showed that the ratio was significantly greater than zero at all frequencies (one-sample Wilcoxon signed-rank test, N = 11, P < 0.003 for all), except 5 Hz (P = 0.06). In addition, the magnitude of this ratio differed significantly across frequencies (linear mixed-effect models, N = 11, P < 0.001), but this frequency-dependence did not significantly vary with rTMS intensity (P = 0.443), indicating a frequency-selective transmission profile that was consistent across stimulation intensities. Contrast analysis revealed significantly greater transmission within the 20-50 Hz range than within both the 5-10 Hz (P = 0.002) and 60-80 Hz ranges (P < 0.001) (**Fig. 4C**). Transmission within the 60-80 Hz range was also lower than within the 5-10 Hz range (P = 0.034). Together, these results demonstrate a frequency-selective, bandpass-like profile of corticospinal transmission during rTMS.

We then evaluated whether the rTMS frequencies transmitted to the motor neuron pool altered the shared synaptic oscillations across spinal motor neurons. To this end, we compared motor unit pooled z-coherence before and during rTMS. **Fig. 5A** shows representative examples for 5, 20, and 50 Hz stimulation. At 5 Hz, z-coherence did not show a clear increase at the stimulation frequency (green rectangle). In contrast, at 20 Hz and 50 Hz, z-coherence at the stimulation frequency was markedly higher during rTMS than before rTMS.

Group analysis confirmed this pattern with the area under the curve of motor unit pooled z-coherence significantly increasing from pre-to during-rTMS for all stimulation frequencies (linear mixed-effect models, N = 7 with ≥ 4 matched motor units, P < 0.025 for all), except 5 Hz (P = 0.099) (**Fig. 5B**). Contrast analysis revealed that increases in common oscillations between motor unit spike trains were significantly greater for 20 Hz than for 5 Hz (P = 0.005) and 10 Hz (P = 0.015).

### E. Simulations Suggest that Excitatory and Inhibitory Post-synaptic Potentials Underlie Bandpass Behavior

To explore potential mechanisms for the bandpass filtering observed experimentally, we simulated the response of a motor neuron pool to rTMS inputs using the motor neuron model proposed by Fuglevand, et al. [44]. We simulated three synaptic input scenarios (**Fig. 6A**). In the first scenario, the common synaptic input was modeled as a train of EPSPs only. In the second scenario, the input consisted exclusively of IPSPs. In the third scenario, each rTMS volley evoked a combination of EPSPs and IPSPs arriving at the motor neuron pool. Simulations revealed that bandpass filtering of corticospinal transmission emerged only when excitatory and inhibitory post-synaptic potentials co-occurred. Simulations with EPSPs only or IPSPs only failed to reproduce the frequency-selective profile observed experimentally (**Fig. S1**). In contrast, the combined EPSP-IPSP scenario produced input-output coupling profiles consistent with a bandpass filter (**Fig. S2**). Within this scenario, we further examined which parameter combinations best reproduced the experimentally observed spectral input-output coupling profile. The closest match to the experimental data was obtained when EPSPs were modeled with τ = 4 ms and IPSPs with τ = 6 ms, with equal synaptic gain for excitatory and inhibitory inputs. Under these conditions, the simulated input-output coupling profile closely resembled the experimentally estimated bandpass profile (**Fig. 6B**). These results provide a potential mechanistic explanation for the frequency-selective transmission observed experimentally and provide quantitative estimates of the underlying post-synaptic potential dynamics.

## IV. DISCUSSION

In this study, we combined HDsEMG decomposition with rTMS to investigate how rhythmic cortical stimulation of different frequencies and intensities are transmitted to spinal motor neurons. Our main findings revealed that, except at 5 Hz, rTMS-induced oscillations are conveyed to the motor neuron pool in a manner consistent with a linear system, with stronger coupling observed at higher stimulation intensities. In addition, corticospinal transmission exhibited marked frequency selectivity, with frequencies within the 20-50 Hz range being preferentially transmitted to the neural drive. This selective transmission was associated with the emergence of oscillatory components in the shared synaptic input that were not present before stimulation. Supported by computational modeling, these findings suggest that frequency-dependent rTMS transmission may arise from the interaction between excitatory and inhibitory post-synaptic potentials and allowed us to estimate potential synaptic dynamics underlying this process. Together, our results provide evidence that corticospinal pathways operate as a frequency-selective linear system, shaping how descending neural inputs are transformed into motor output.

### A. Linearity of Corticospinal Transmission During rTMS

The transformation of synaptic input into motor neuron discharge at the level of single motor neurons is inherently nonlinear, due to their threshold-dependent firing and intrinsic membrane properties, such as persistent inward currents [20, 69–71]. This nonlinearity is further exacerbated by the relatively low discharge rates of spinal motor neurons during low to moderate contraction levels [72], which can lead to undersampling of the input signal, particularly at higher frequencies (for a review, see Farina and Negro [73]). This limitation can also be interpreted in terms of sampling theory, where frequencies higher than approximately half of the firing rate cannot be faithfully represented [74]. However, theoretical, computational, and experimental evidence has demonstrated that when a small number of motor neurons receive a common synaptic input, the collective output of the motor neuron pool provides a more faithful representation of the input than that of individual neurons [20, 45, 70, 75]. In this case, the effective sampling of the input is no longer limited by the discharge rate of individual neurons but by the aggregate density of spikes across the population [76]. In other words, the motor neuron pool effectively linearizes the input-output relationship, a phenomenon also observed in other neuronal systems [77, 78]. Consistent with this, common synaptic input has been identified as the primary determinant of the neural drive to muscle [56, 75, 79, 80].

The results of the present study support this view. During low-force contractions (10% MVC), rTMS-induced oscillatory inputs were reflected in the discharge activity of the majority of motor neurons, independently of their individual discharge rates, which ranged from ∼5-20 pps (**Fig. 2B**-**D**). Not all individual motor neurons exhibited significant coupling with the input (**Fig. 2C**), which is consistent with their intrinsic nonlinear properties and limited ability to faithfully encode synaptic inputs. This effect became even more evident when considering the cumulative activity of the motor neuron pool, where rTMS inputs were strongly coupled with oscillations in the neural drive (**Fig. 3**). Furthermore, input-output coupling scaled proportionally with subthreshold rTMS intensity (**Fig. 4A**-**B**), consistent with an approximately linear transmission at the motor neuron population level. These results were consistent across stimulation frequencies above 5 Hz, suggesting that approximate linear transmission emerges at the level of the motor neuron pool within a specific frequency range. These findings align with previous observations showing an approximately linear relationship between TMS intensity and MEP amplitude when background force is held constant [81], although it is important to note that across a broader range of TMS intensities and activation levels, this relationship can become sigmoidal [36, 37]. Taken together with previous studies, our findings provide direct evidence that corticospinal inputs are widely distributed across the motor neuron pool and support the view that population level motor neuron activity enables a near-linear transformation of descending synaptic inputs into motor output.

### B. Frequency-selective Corticospinal Transmission during rTMS Arises from Synaptic Interactions

Corticospinal transmission during rTMS exhibited a clear frequency-selective behavior, with oscillatory inputs within 20-50 Hz being preferentially transmitted to the spinal motor neuron pool (**Fig. 4C**). In contrast, low-frequency stimulation at 5 Hz did not result in significant transmission, whereas higher-frequency components above 60 Hz were markedly attenuated. These findings indicate that corticospinal pathways behave as a bandpass system, selectively transmitting oscillatory rTMS inputs within a specific frequency range to the neural drive.

Frequency-dependent filtering of synaptic inputs has been previously reported in both cortical and somatosensory neurons in rats, where intrinsic membrane and synaptic properties shape the spectral content of neuronal output [82, 83]. In the present study, the absence of transmission at low frequencies and attenuation at higher frequencies suggests that corticospinal pathways do not simply relay cortical inputs during rTMS, but actively transform them through frequency-dependent mechanisms. While the precise origin of these filtering properties cannot be directly determined from the experimental data, the interaction between rTMS-induced activity and the ongoing voluntary drive may contribute to the reduced transmission of low-frequency inputs. During the 10% MVC contraction, motor neurons were already active, receiving continuous synaptic input and discharging at low to moderate rates [72]. Under these conditions, low-frequency rTMS pulses may act as isolated perturbations superimposed on ongoing input and, thus, may not impose a consistent oscillatory modulation across the motor neuron pool. In contrast, higher-frequency stimulation provides an input capable of interacting with the ongoing synaptic drive and entraining motor neuron activity, leading to stronger input-output coupling even with low intensity stimulation. Additionally, the attenuation of low-frequency inputs may reflect active network mechanisms, such as spinal interneuron circuits or recurrent inhibition, which have been suggested to suppress slow oscillations in corticospinal pathways [26, 27, 84]. Finally, although stimulation duration was matched across frequencies, the total number of pulses delivered and their temporal distribution differed, particularly between the continuous 5 Hz stimulation and the higher-frequency protocols delivered in blocks. Therefore, we cannot exclude the possibility that these methodological differences contributed to the weaker corticospinal coupling observed at 5 Hz. Importantly, our findings are consistent with previous observations showing that spinal motor neurons transmit coherent oscillations within a limited bandwidth, typically up to ∼70-80 Hz [22–25].

To further investigate the mechanisms underlying this bandpass behavior, we simulated the response of a motor neuron pool to rTMS-like inputs under different synaptic configurations. Notably, only when synaptic inputs were modeled as a combination of excitatory and inhibitory post-synaptic potentials (EPSP-IPSP scenario; **Fig. 6**), the system reproduced the bandpass filtering observed experimentally. This finding is consistent with evidence that the net effect of rTMS is a complex interplay between the recruitment of excitatory and inhibitory circuits [48, 85]. From a signal processing perspective, the combination of excitatory and inhibitory synaptic inputs (i.e., biphasic response) naturally gives rise to bandpass filtering properties [86]. Importantly, specific combinations of synaptic dynamics were required to match the experimental observations. The bandpass characteristics were best reproduced when EPSPs were modeled with a time constant of 4 ms and IPSPs with a time constant of 6 ms, with comparable synaptic gains. These results suggest that the temporal interplay between excitation and inhibition critically shapes the frequency-dependent transmission of corticospinal inputs, consistent with general neural coding principles where dynamic inhibition expands transmission bandwidth [86]. Together, these findings provide a mechanistic framework for understanding how corticospinal pathways filter oscillatory inputs and offer novel insight into the synaptic dynamics underlying rTMS-induced neural activity.

### C. Implications for Neuromodulation of Corticospinal Pathways

Our results bridge the gap between fundamental principles of neural control of human movement and their potential clinical applications, providing insight into how corticospinal inputs can be effectively modulated. Specifically, rTMS delivered at frequencies between 10 and 60 Hz, and particularly at 20 Hz and 50 Hz, resulted in the strongest input-output coupling, indicating that these stimulation parameters are more effectively transmitted to spinal motor neurons. These observations suggest that corticospinal pathways exhibit preferential sensitivity to specific frequency ranges, which may be leveraged to enhance neuromodulatory interventions aimed at increasing corticospinal excitability. This interpretation aligns with previous work using TMS paradigms showing that targeted modulation of corticospinal transmission, specifically through timing-dependent plasticity mechanisms at cortico-motoneuronal synapses, can enhance voluntary motor output and promote recovery [30, 31, 87]. This interpretation is also consistent with previous work demonstrating frequency-dependent tuning of the human motor system, including studies using transcranial alternating current stimulation that identified a “natural frequency” of the motor cortex around the beta band (∼20 Hz) [88], as well as with theta-burst stimulation protocols that incorporate high-frequency bursts (e.g., 50 Hz) to induce lasting changes in corticospinal excitability [85].

Interestingly, rTMS delivered within this frequency range (20-50 Hz) induced oscillatory components in motor unit activity at the stimulation frequency that were not present before stimulation (**Fig. 6**), suggesting a direct modulation of the shared synaptic input to the motor neuron pool. From a methodological perspective, these changes were not detected by conventional bipolar EMG amplitude (**Table S1**), which are widely used in both research and clinical settings. This finding highlights the added value of HDsEMG decomposition in revealing neural control mechanisms during rTMS that are not captured at the global EMG amplitude level. In this context, combining non-invasive stimulation techniques with high-resolution motor unit analysis may provide a powerful framework to probe how corticospinal inputs are transformed into motor output at the level of the motor neuron pool. Together, these results not only provide mechanistic insight into corticospinal transmission but also highlight the importance of considering both stimulation parameters and measurement techniques when designing and interpreting neuromodulation protocols.

Several limitations of the present study should be acknowledged. First, stimulation intensities were limited to a maximum of 70% RMT, as pilot testing indicated that higher intensities could evoke overt motor responses and interfere with the steady contraction. Accordingly, no significant differences in force steadiness were observed between pre-and during-rTMS across the tested conditions (**Table S1**). Second, the experiments were performed at a relatively low contraction level (10% MVC) to allow accurate decomposition and tracking of motor units. As motor neuron discharge rates and synaptic integration properties vary with contraction intensity [22, 72], it remains unclear how increasing force levels may influence the transmission of rTMS inputs through corticospinal pathways. Future work should examine how corticospinal frequency transmission interacts with different levels of voluntary drive and different physiological modifiers (age, sex, comorbidities). Moreover, the paradigm could be applied to different pathological conditions impacting on cortico-spinal motor control, including primary first vs second motor neuron pathology, amyotrophic lateral sclerosis or basal ganglia diseases, known to be associated with altered neurophysiological motor control [89]. Third, the present findings were obtained from a single muscle group (thenar muscles) and, although this model provides a well-controlled framework to study corticospinal transmission, the extent to which these results generalize to other muscles, particularly proximal or lower-limb muscles, remains to be determined. Finally, the computational simulations used a simplified representation of TMS-evoked synaptic inputs to spinal motor neurons. Although individual TMS pulses may generate multiple descending volleys through complex cortical interactions, as demonstrated in more detailed computational models [90], these responses were approximated by a single effective postsynaptic response per stimulation pulse. Consequently, the model does not account for the precise timing and relative contributions of individual descending volleys or the cortical dynamics underlying their generation. Therefore, although the simulations demonstrate that combined excitatory and inhibitory postsynaptic inputs can reproduce the experimentally observed frequency selectivity, they do not establish that this is the only mechanism responsible for the bandpass-like transmission.

