## Supplementary material for "Bandpass corticospinal transmission during repetitive TMS revealed by motor unit recordings": Table S1; Fig. S1; Fig. S2

Hélio V. Cabral *et al.*

**This PDF file includes:**

Table S1

Figs. S1 to S2

**Table 1. Force steadiness and bipolar EMG amplitude before and during rTMS.** Values are mean ± standard deviation across participants for each rTMS frequency and stimulation intensity (% resting motor threshold, RMT). Force steadiness is expressed as the coefficient of variation of force. EMG root-mean-square (RMS) amplitude was normalized to the RMS amplitude recorded during maximal voluntary contraction (MVC).

| **rTMS frequency (Hz)** | **rTMS intensity (%RMT)** | **Coefficient of variation of force (%)** | | **Bipolar EMG RMS amplitude (%MVC)** | |
|---|---|---|---|---|---|
| | | Pre-rTMS | During-rTMS | Pre-rTMS | During-rTMS |
| 5 | 50 | 4.33 ± 2.54 | 3.37 ± 1.48 | 12.81 ± 7.46 | 13.58 ± 7.89 |
| | 60 | 3.37 ± 0.58 | 3.37 ± 1.12 | 12.96 ± 9.58 | 14.84 ± 10.72 |
| | 70 | 3.77 ± 1.51 | 3.74 ± 1.31 | 13.37 ± 8.33 | 16.81 ± 10.04 |
| 10 | 50 | 3.15 ± 1.20 | 2.96 ± 1.10 | 14.14 ± 8.49 | 15.28 ± 9.42 |
| | 60 | 4.20 ± 1.74 | 3.85 ± 1.75 | 16.24 ± 8.55 | 17.21 ± 9.37 |
| | 70 | 3.21 ± 1.32 | 3.41 ± 1.22 | 13.52 ± 8.90 | 15.66 ± 11.83 |
| 20 | 50 | 5.18 ± 2.96 | 3.72 ± 1.12 | 14.75 ± 8.23 | 16.92 ± 10.86 |
| | 60 | 3.32 ± 1.04 | 3.24 ± 1.57 | 13.21 ± 8.72 | 15.74 ± 11.38 |
| | 70 | 3.36 ± 1.23 | 3.78 ± 1.87 | 12.46 ± 10.80 | 15.67 ± 12.50 |
| 30 | 50 | 3.34 ± 0.87 | 2.85 ± 0.93 | 11.03 ± 7.21 | 12.90 ± 9.30 |
| | 60 | 4.12 ± 1.93 | 3.73 ± 1.23 | 12.52 ± 7.77 | 14.23 ± 8.83 |
| | 70 | 3.85 ± 1.55 | 4.36 ± 1.48 | 16.23 ± 12.59 | 20.27 ± 15.17 |
| 50 | 50 | 3.77 ± 0.78 | 3.40 ± 1.29 | 17.94 ± 12.46 | 19.45 ± 13.65 |
| | 60 | 3.61 ± 1.91 | 3.14 ± 1.49 | 14.72 ± 12.34 | 17.37 ± 14.89 |
| | 70 | 4.15 ± 1.42 | 4.31 ± 2.68 | 11.87 ± 7.79 | 17.46 ± 13.40 |

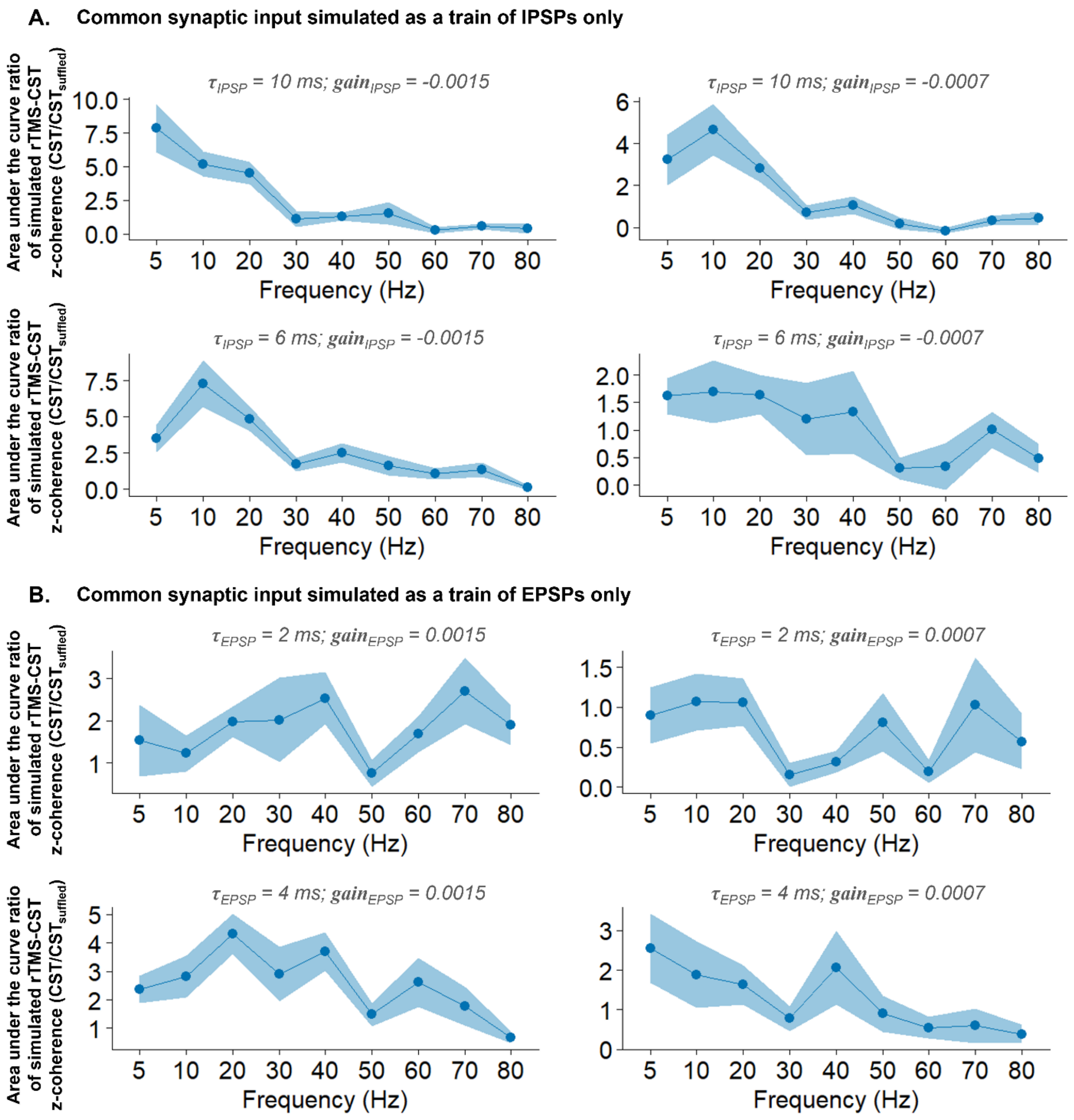


**Fig. S1. Simulated corticospinal transfer functions with purely inhibitory or excitatory synaptic inputs.** Transfer functions obtained when the common synaptic input to the motor neuron pool was modeled as a train of inhibitory post-synaptic potentials (IPSPs) only (A) and as a train of excitatory post-synaptic potentials (EPSPs) only (B). Four parameter combinations are shown for each case, with varying the IPSP time constant (τ = 6 or 10 ms), the EPSP time constant (τ = 2 or 4 ms) and synaptic gain. For each condition, the transfer function is expressed as the area under the curve ratio of rTMS-CST z-coherence relative to the shuffled CST across frequencies (5–80 Hz).

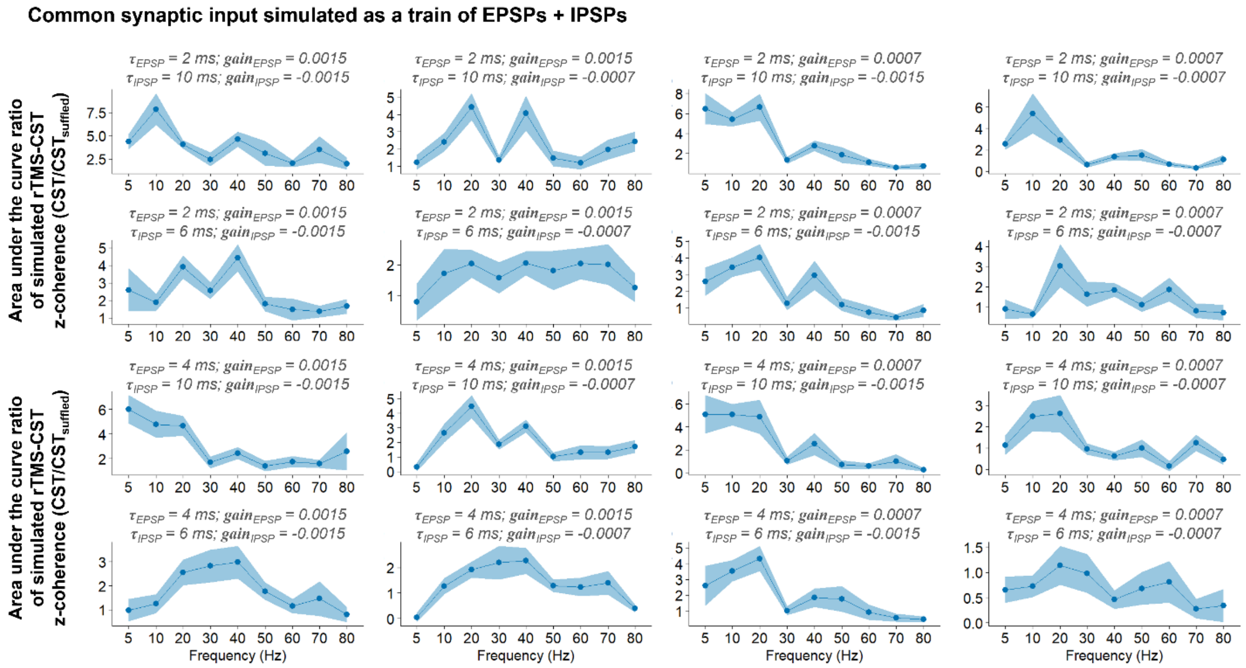


**Fig. S2. Simulated corticospinal transfer functions with combined excitatory and inhibitory synaptic inputs.** Transfer functions obtained when the common synaptic input to the motor neuron pool was modeled as a train of combined excitatory and inhibitory post-synaptic potentials (EPSP + IPSP). Different parameter combinations are shown, varying EPSP time constants ($\tau_{EPSP}$ = 2 or 4 ms), IPSP time constants ($\tau_{IPSP}$ = 6 or 10 ms), and synaptic gains. For each condition, the transfer function is expressed as the area under the curve ratio of rTMS-CST z-coherence relative to the shuffled CST across frequencies (5–80 Hz). The combination of excitatory and inhibitory inputs with specific synaptic dynamics (bottom left panel) reproduced a bandpass profile, with enhanced transmission in the 10–60 Hz range, consistent with the experimental observations
